# The ϕPA3 Phage Nucleus is Enclosed by a Self-Assembling, 2D Crystalline Lattice

**DOI:** 10.1101/2022.04.06.487387

**Authors:** ES Nieweglowska, AF Brilot, M Méndez-Moran, M Baek, J Li, C Kokontis, Y Cheng, D Baker, J Bondy-Denomy, DA Agard

**Affiliations:** Department of Biochemistry, University of California San Francisco, San Francisco, CA 94143; Sauer Structural Biology Laboratory, Center for Biomedical Research Support, University of Texas at Austin, Austin, Texas; Department of Biochemistry, University of Washington, Seattle, WA 98195; Institute for Protein Design, University of Washington, Seattle, WA 98195; Department of Microbiology, University of California San Francisco, San Francisco, CA 94143; Howard Hughes Medical Institute; Howard Hughes Medical Institute, University of Washington, Seattle, WA 98195

## Abstract

A growing number of jumbo bacteriophages, with genomes exceeding 200 kb, have been found to establish a Phage Nucleus—a micron-scale, proteinaceous structure encompassing the replicating phage DNA. Bacteriophage and host proteins associated with replication and transcription are concentrated inside the Phage Nucleus while nucleotide synthesis, translation, and numerous other host and exogenous proteins are effectively excluded, including CRISPR-Cas and restriction endonuclease host defense systems. Here, we show that fragments of the Phage Nucleus isolated from ϕPA3 infected *Pseudomonas aeruginosa* cells form a square lattice and demonstrate that the recombinantly purified primary *Ph*age N*u*clear E*n*closure (PhuN) protein spontaneously assembles into sheets also constructed from a square lattice which we resolve to 3.8 Å by cryo-EM. Our structure reveals that the flexible termini and large loops mediate adaptable inter-tetramer contacts that drive shell assembly into a C2-symmetric lattice. While the interfaces between subunits are mostly well packed, two of the interfaces are open, forming clear channels that likely have important functional implications for the transport of proteins, mRNA, and small molecules.

## Introduction

There is a constant evolutionary pressure for bacteria to develop defense mechanisms against invading bacteriophages and for the phages to develop effective countermeasures.^1^ To that end, restriction-modification and numerous CRISPR systems are widespread amongst bacterial hosts while phages have developed their own DNA modification and anti-CRISPR systems.^2^ A subset of so-called jumbo bacteriophages, defined by having genomes exceeding 200 kb, have recently been shown to encode an elaborate system for sequestering the phage genome away from host nucleolytic attack, conveying broad resistance to targeting by the host.^3^ This is accomplished via the assembly of a selectively permeable protein shell that encompasses the phage genome.^4,5^ The shell with its contents is referred to as the Phage Nucleus for its remarkable functional similarity to the eukaryotic nucleus: this structure forms a selectively permeable compartment that is centered within the host bacteria by a dynamic bipolar spindle formed from a phage encoded, divergent tubulin called PhuZ.^4,6–8^ The bacteriophage and host proteins involved in phage replication and transcription are concentrated within the Phage Nucleus shell while translation and nucleotide synthesis machinery, the aforementioned CRISPR-Cas and restriction endonucleases, as well as other host or exogenous proteins are effectively excluded.^3,4^ Unlike other proteinaceous prokaryotic compartments, such as carboxysomes or viral capsids, this shell can grow significantly throughout infection to reach nearly micron-scale, a process likely driven by genome replication.^4,9–11^ In contrast to some other membraneless compartment forming systems, such as protein condensates, subunits do not diffuse throughout the shell of the Phage Nucleus, indicative of unique assembly properties.^4,12^

While the total protein composition of the Phage Nucleus shell is unknown for any jumbo phage, Gp105 from the ϕKZ family phage 201ϕ2-1 was shown to be a marker for the shell.^4^ It is the most highly expressed protein that is not part of the mature phage particle.^4^ Here we formally introduce this protein and its related family members as *ph*age n*u*cleus e*n*closure or PhuN proteins. The PhuN family currently includes Gp53 (ϕPA3), Gp54 (ϕKZ), Gp105 (201ϕ2-1), Gp202 (PCH45), and numerous putative homologous proteins from newly sequenced jumbo bacteriophages.^4,5,13,14^ Beyond these other phage proteins, PhuN has no clear previously characterized homologous relatives. This unique assortment of biophysical and biological properties combined with the mystery about the tertiary structure of PhuN established it as an exciting target for structural and biochemical analysis.

In this work, we demonstrate that Phage Nucleus fragments isolated from ϕPA3 infected *P. aeruginosa* cells form a square lattice. We further show that Gp53, the PhuN family member from bacteriophage ϕPA3, readily assembles into large 2D lattices *in vitro*. We utilized a limited tilt, cryo-electron microscopy data collection scheme paired with single particle processing to determine a 3.8 Å map of PhuN assemblies. Assisted by RoseTTAFold^15^ and AlphaFold,^16^ we present a model and analysis of the assembly enclosing the Phage Nucleus. The structure reveals that the flexible termini and large loops mediate adaptable inter-tetramer contacts that drive shell assembly into a C2 symmetric lattice. While the interfaces between subunits are mostly well packed, two of the interfaces are open, forming clear channels that likely have important functional implications. Despite the lack of detectable sequence homology, analysis of the atomic structure reveals that the basic architecture is derived from a fusion of acetyltransferase and carbon monoxide dehydrogenases/nuclease domains.

## Results

### Isolation of Phage Nucleus Fragments

*P. aeruginosa* expressing maltose-binding protein (MBP)-PhuN fusions were infected with ϕPA3 and lysed after 60 minutes of infection. While the MBP tag was intended to aid affinity purification, differential centrifugation of the lysates proved more successful at shell isolation, likely due to the large assemblies hindering efficient interaction with amylose beads. The resulting isolates resemble fragmented shells and have a clearly defined lattice structure **(Fig.1a)**. Two-dimensional averaging of the isolated fragments further reveals a lattice with clear deviations from C4 symmetry but not enough detail to assign C2 or C1 symmetry **(Fig.1a)**. The subunits forming the lattice appear to be a single, approximately 60 by 40 Å protein species. Recent low resolution *in situ* tomography of Phage Nuclear Shells from phages 201ϕ2-1 and Goslar reports C4 symmetry but is otherwise consistent with this observation, demonstrating a square lattice with similarly sized subunits.^17^ The isolated shell fragments often appear folded over and the lattice can vary in directionality, suggestive of both structural plasticity and imperfections or locally altered symmetry within the lattice of endogenous mature Phage Nuclear Shells **(Fig. 1a; S1a,b)**.

**Fig. 1.**
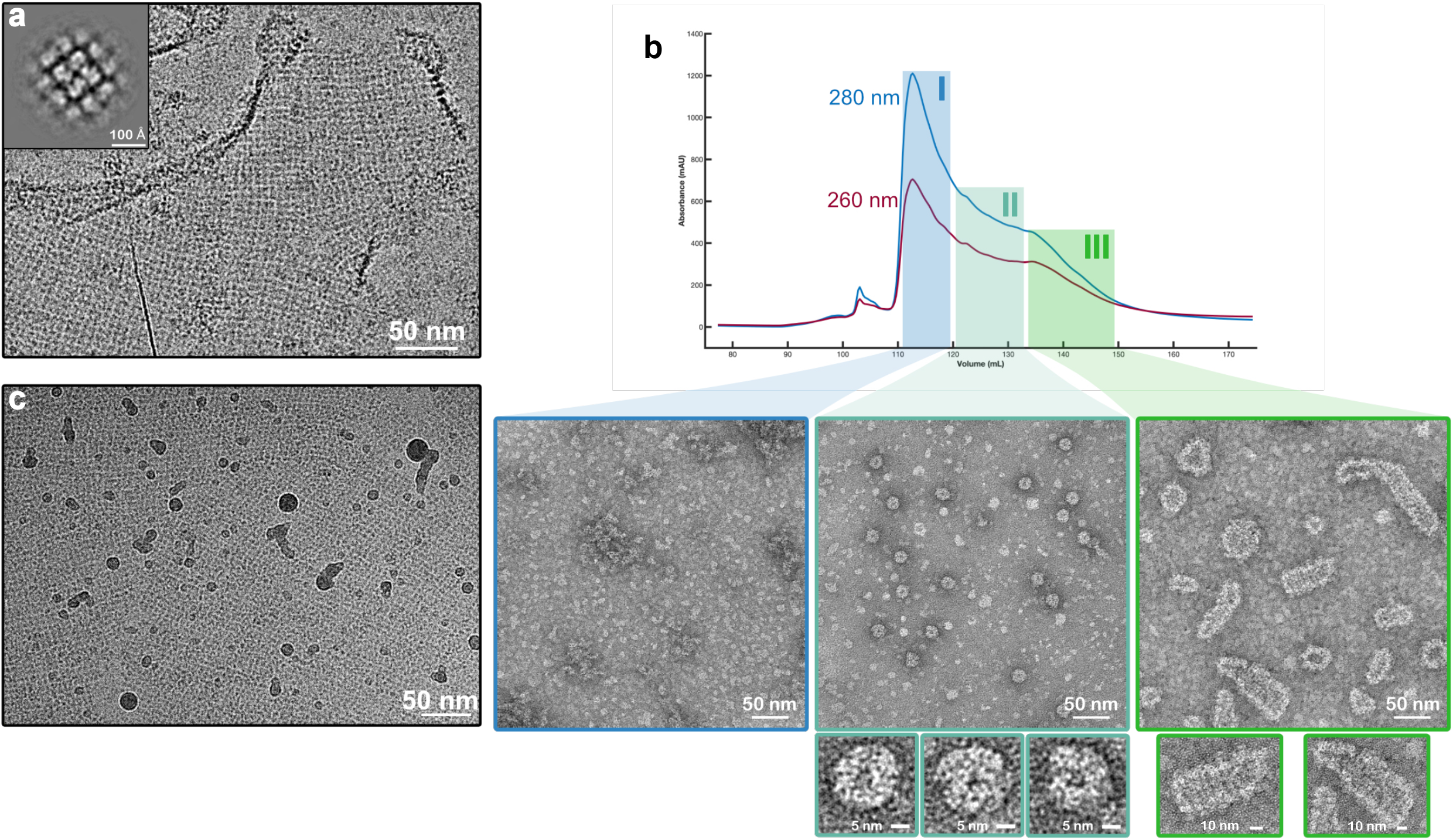
PhuN forms broad range of assemblies. **a**, Phage Nuclear Shell fragments isolated from ϕPA3 infected *Pseudomonas aeruginosa*. Folds and differences in lattice orientation can be observed. The inset shows a 2D class average from processing the fragments. The symmetry does not appears C2 or C1. **b**, Anion exchange trace showing extended elution profile and corresponding negative stain EM micrographs. Class I (blue) shows concentrated monomeric species. Class II (turquoise) shows medium sized assemblies in the 25 nm range. Class III (green) reveals assemblies with elongated shapes and striated texture. **c**, *in vitro* assembled 2D crystalline arrays at 0° tilt showing three main patches with distinct lattice orientations and gentle curvature. Boarders between different crystal patch orientations appear smooth.

### PhuN Self-Assembles During Purification

ϕPA3 PhuN is a 66.6 kDa protein with marginal solubility upon expression in *E. coli*, a problem surmounted by the addition of an N-terminal maltose-binding protein (MBP) solubility tag. Importantly, the addition of MBP does not prevent PhuN from incorporating into the Phage Nuclear Shell during ϕPA3 infection in *P. aeruginosa* **(Fig. S1c)**. All *in vitro* data presented here utilize the MBP-PhuN fusion and, for simplicity, will be referred to as PhuN unless otherwise specified.

During purification, exposure to an anion exchange column induced the formation of a broad spectrum of PhuN assemblies observed upon elution **(Fig. 1b)**. While the exact reason for this preferential on-column assembly is not known, it likely occurs as a result of increased local PhuN concentration during sample application, binding, and elution. Exposure to the high surface charge density of the column resin may also play a role.

These assemblies were grouped into three broad classes based on their appearance in negative stain election microscopy. Class I primarily contains monomeric PhuN **(Fig. 1b)**. Class II includes rounder species approximately 25 nm in diameter with distinct edges. Despite efforts to optimize the production of these for structure analysis, they proved to be an elusive species, often outnumbered by more irregular assemblies like those found in Class III **(Fig. 1b)**. Class III consists of highly variable 50-130 nm long, elongated assemblies **(Fig. 1b)**. The Class III species revealed a striated organization highlighting the striking diversity in both the shape and size of PhuN assemblies. In cryoEM, although heterogeneously shaped, Class III assemblies appear to have a more regular, lattice-like organization and showed a density around the exterior consistent with the presence of the N-terminally fused MBP **(Fig S1d, top)**.

### Growing and Imaging PhuN 2D Lattices

We initially set out to determine conditions that would allow us to control the assembly rate and state of PhuN. Using the anion exchange column as an assay to monitor shifts in assembly states, we instead found that at a pH of 6.5 monomeric PhuN forms large, 2D polycrystalline assemblies when applied to negatively charged grids for negative stain EM **(Fig S2b)**. The low pH was also necessary to maintain the integrity of shell fragments isolated from infected cells, suggesting that pH may be an important factor to consider in shell assembly *in vivo*. Stress induced by antibiotic exposure has been observed to decrease the cytosolic pH to 6.7 in *P. aeruginosa* and other bacteria.^18^ Perhaps phage infection may act as a similar metabolic stressor and have an analogous impact on pH.

These 2-dimensional crystals were further optimized using a 2D surface crystallization method as often used in 2D electron crystallography.^19^ Purified His-tagged PhuN was diluted into a buffer droplet in Teflon wells that had a pre-formed lipid monolayer containing 21% Ni-NTA modified lipids **(Fig. S3)**. This approach yielded larger, better ordered crystalline assemblies than both those observed in negative stain and those isolated from infected cells. These arrays were then prepared for and imaged by cryo-EM. Despite optimizing lattice formation, the 2D sheets generally included many local regions of distinct lattice orientation as well as numerous bends and waves **(Fig. 1c, S4)**. Consequently, we chose a predominantly real-space approach, analogous to single particle analysis,^20^ instead of the more common 2D electron crystallography strategy **(Fig. S5)**. To obtain the necessary views, data were collected at multiple tilt angles: 0°, 15°, 30°, 35°, 40°, 45°, 50°, 60° (**Fig. S4)**. In order to compensate for the high degree of beam induced motion and lower image quality resulting from increased sample thickness in our tilted samples, data were collected at a high frame rate and processed using a tilt-optimized version of MotionCor2.^21^ On conventional grids, few lattice patches could be found and those that did adsorb were only visible in very thick ice. We overcame this challenge by utilizing amino functionalized graphene oxide grid supports ^22^ for the bulk of our data acquisition as they stabilized the lattices in somewhat thinner ice and greatly increased the number of lattice patches adsorbed. Using this approach, we were able to resolve PhuN assemblies in 2D arrays to ∼3.8 Å and build an atomic model starting with predictions from RoseTTAFold^15^ and AlphaFold^16^ **(Fig. 2b-d, S5)**.

**Fig. 2.**
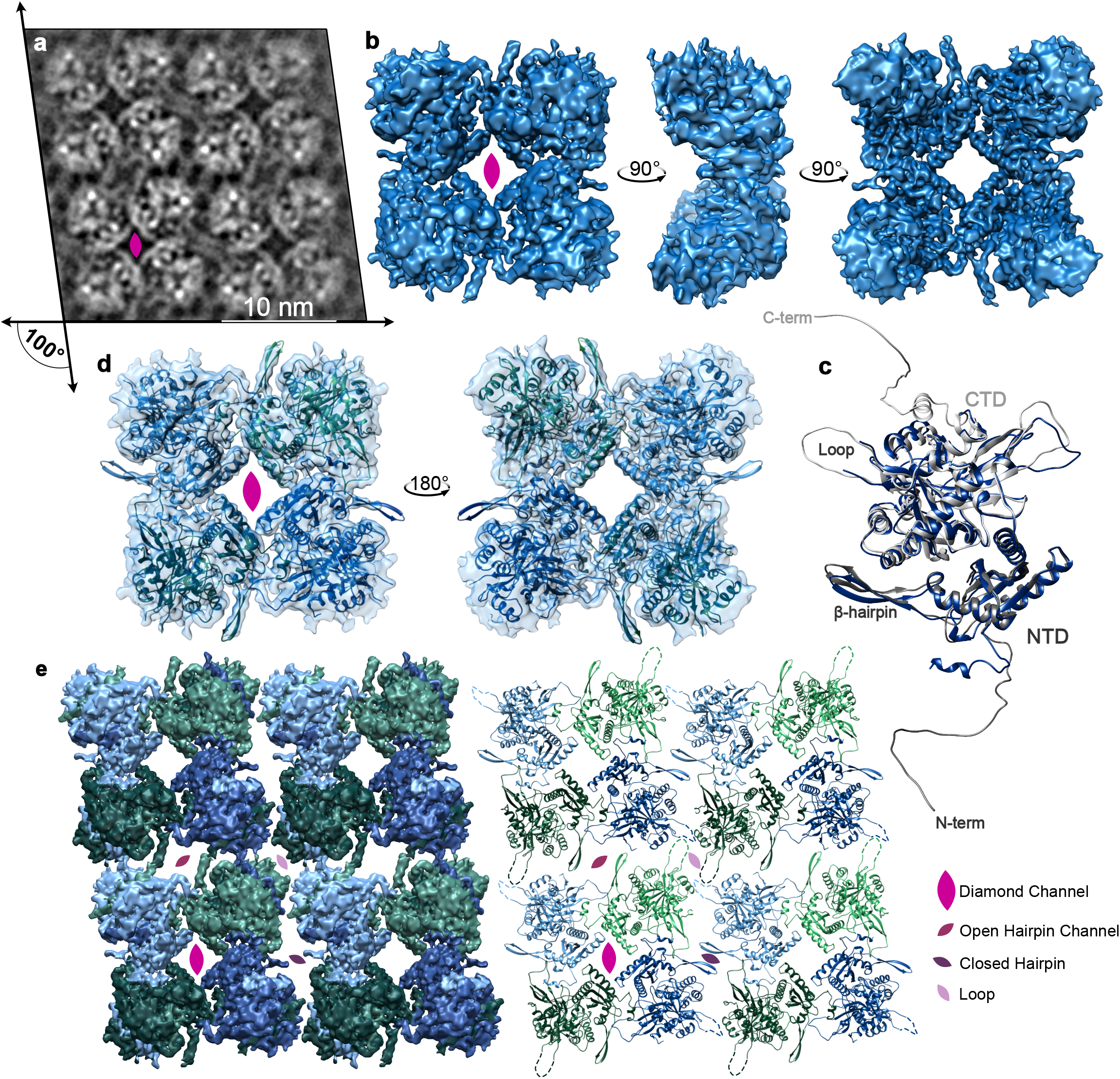
PhuN assembles into a C2 Symmetric Lattice. **a**, Example 2D class average highlighting the C2 symmetry and resulting four unique interfaces. The lattice is skewed to an approximately 100° angle. The positioning of the flexible loops that are difficult to resolve in 3D is visible. **b**, Highest resolution map of the core tetramer centered on the Diamond interface as viewed from different angles. **c**, The final Phun-C subunit model (blue) overlaid with the AlphaFold prediction used as our starting model (grey). Residues 1-18, 276-287, 556-602 had no corresponding density in our map and were deleted from the model. **d**, The tetrameric model of PhuN-C and PhuN-O fit into the map in b. **e**, The map from b segmented and positioned to recreate the channels observed in 2D classes (left) paired alongside the corresponding 16mer-model (right). In both the segmented maps and models, PhuN-O is represented in blue while PhuN-C subunits are represented in green. Unresolved and unmodeled loops are represented with dashed lines to reflect the 2D class while the C-terminal tails are excluded.

### PhuN Lattices: 4 Unique Interfaces

The ϕPA3 PhuN monomer is comprised of two connected domains. Despite the lack of convincing sequence homology to other proteins, structural comparison of our model with coordinates in the PDB using the Dali Structure Comparison Server^23^ revealed that the large domain is structurally homologous to acetyltransferases while the smaller domain has a less clear fold homology, sharing similarities with proteins including carbon monoxide dehydrogenases/Acetyl-CoA Synthases as well as several restriction endonucleases (**Fig. S10)**. As other proteins forming nanocompartments or encapsulins often share structural similarities with capsids,^24^ the PhuN homologies suggest a unique evolutionary origin. As no related binding sites or catalytic residues could be identified in PhuN, it is unlikely that either domain has retained any ancestral catalytic activity. It remains to be seen whether these homologies have functional implications beyond a purely structural role.

PhuN assembles into a tetrameric lattice with a dimeric asymmetric unit, giving rise to C2 symmetry **(Fig 2a)**. Alignment of the asymmetric subunit models using the acetyltransferase-like domain results in an RMSD of ∼2 Å. The bulk of the differences between the structured domains of asymmetric subunits appear to arise from small shifts in various helices as well as interconnecting loops. Each core tetramer is held together by tight packing of four monomers with both domains contributing, while interactions between the tetramers are looser and predominantly coupled via highly flexible elements (loops, termini, etc.). In keeping with the C2 symmetry, each PhuN tetramer forms four unique interfaces, two of which form channels: a Diamond channel at the center of the core tetramer, an Open Hairpin channel and Closed Hairpin interface forming the lateral contacts between core tetramers, and a Loop interface where the outer corners of four tetramers meet **(Fig. 2a,e)**.

The central Diamond channel is lined by the four smaller PhuN domains of the core tetramer. The resulting diamond-shaped opening has a highly negative charge **(Fig. 3b)** and measures 32 Å by 33 Å across the corners. In an elegant domain swap that holds the subunits tightly packed around the channel, a small positively charged helix within the N-terminal 37 residues of the nuclease domain threads into the neighboring subunit, nestling at the top within a negatively charged pocket **(Fig3a)**.

**Fig. 3.**
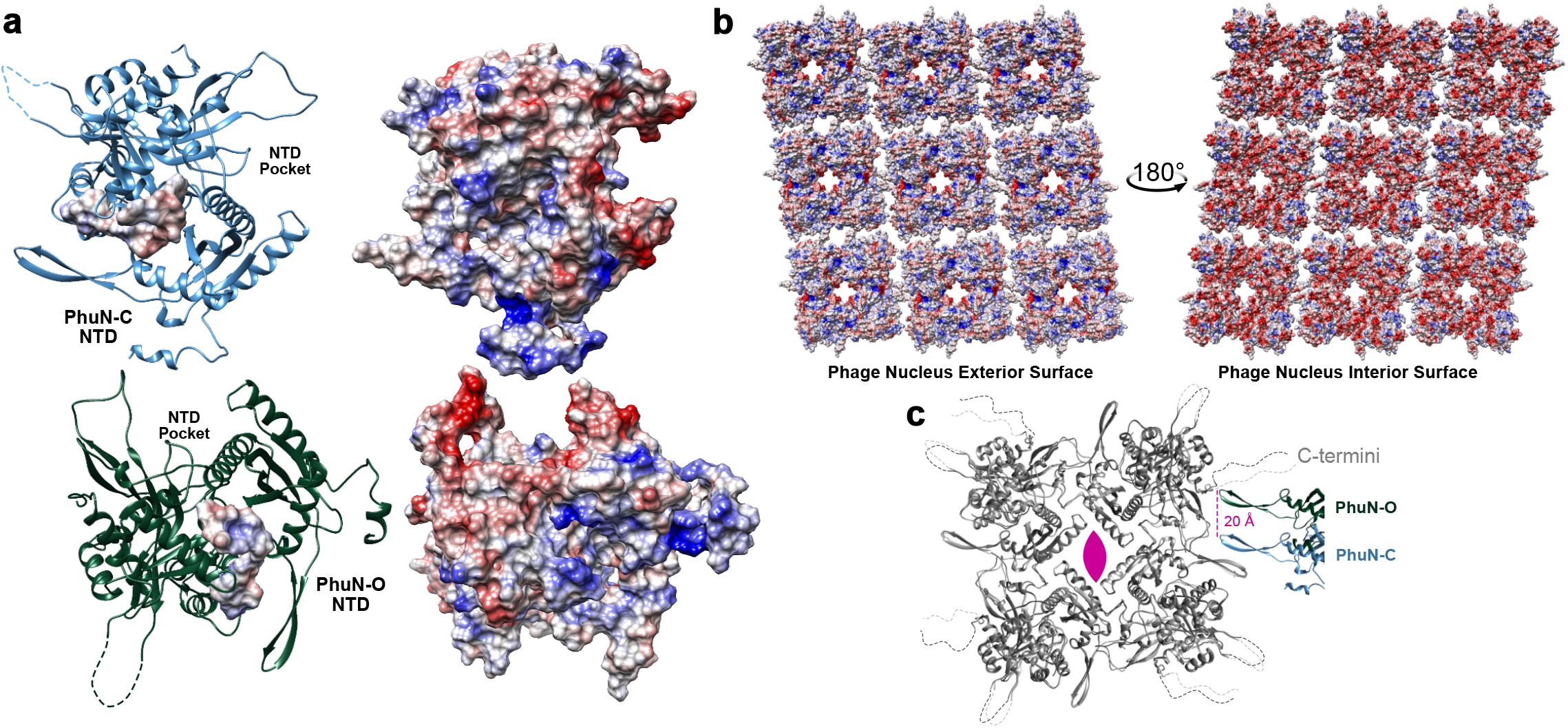
A closer look at PhuN interactions and electrostatic potential. **a**, Surfaces of PhuN-C (top) and Phun-O (bottom) colored by electrostatic potential with their corresponding models on the left. Within the models reside the surfaces of the C-terminal tails colored by electrostatic potential. **b**, Electrostatic potential coloration of the interior and exterior Phage Nucleus surfaces created by PhuN assemblies. **c**, Direct comparison of PhuN-C and PhuN-O. The asymmetric subunits in the tetramer are overlaid showing minor differences (light and dark grey) while the interaction with the neighboring β-hairpins differs dramatically, shifting by approximately 20 Å from near the C-terminus (PhuN-O) towards a flexible loop (PhuN-C).

The Open Hairpin channel and Closed Hairpin interface are located at the lateral interface of two core tetramers. These lateral interactions are established by the same β-hairpins making contacts with different residues of the neighboring tetramers, approximately 20 Å apart **(Fig. 3c, S7a)**. At the Open interface, the tips of opposing β-hairpins (residues 111-126) interact with a positively charged region near where the C-terminus leaves the structured acetyltransferase-like domain of the neighbors **(Fig. S6a,b)**. Keeping these hairpins apart creates a somewhat narrow channel, measuring 31 Å by 41 Å across the corners of the channel. At the Closed interface, these opposing β-hairpins form direct lateral interactions that close the channel and move the tips of the β-hairpins closer to a structured loop (residues 245-258) **(Fig. 3c, S6c)**. As we do not observe any large conformational changes between the asymmetric subunits, this suggests that the 20 Å shift between the β-hairpin positioning at the Open versus Closed Hairpin interfaces arises from a largely rigid motion of the entire core tetramer and thus, as a result, introduces the 10° skew in the lattice network **(Fig 2a; S7b,c)**. One conjecture is that perhaps these are not fixed interfaces, but rather represent Open and Closed states that the lattice can adopt for a functional purpose such as relieving tension in the nearly micron-scale assemblies or adjusting the channel created at the Open interface **(Fig. S7, Movie1)**. Going forward, we will refer to the asymmetric subunits by the positioning of their β-hairpins such that the β-hairpin from PhuN-O forms the Open channel while the PhuN-C β-hairpin forms the Closed interface.

The Loop interface appears to be, in part, held together by interactions between the large flexible loops (residues 275-289) from at least two different subunits. While the full densities of these loops remain unresolved in our 3D map, they are clearly visible in 2D classes **(Fig. 2a)**. The loops from two PhuN-O subunits from opposing core tetramers interact laterally **(Fig. 2a)**. The loops belonging to the PhuN-C subunits at those interfaces are not visible suggesting they are unstructured or in an alternative, less visible conformation in the 2D class. These loops account for the largest conformational differences we observed between asymmetric partners as the core domains of the proteins do not experience sizable conformational changes.

The final 12 residues of the C-terminal tail reside in a groove between the two domains of PhuN, right over the β-hairpin **(Fig. 2e,3a)**. We also see strong densities in our high-resolution map for what is likely the C-terminal strands at the back of the acetyltransferase-like domain **(Fig. 2e)**. These densities differ in the two asymmetric subunits. Unfortunately there was insufficient local resolution to trace how the C-terminus crosses-over between tetramers. This leaves some ambiguity as to which exact paths the long termini take to cross the Open Hairpin channel and Closed Hairpin interface before resting their final 12 residues in the positively charged groove **(Fig. 4)**. An interesting possibility is that these regions may be free and unstructured to allow some flexibility for the tetramers to accommodate alternating between Open and Closed Hairpin states, shell curvature, or different lattice orientations.

**Fig. 4.**
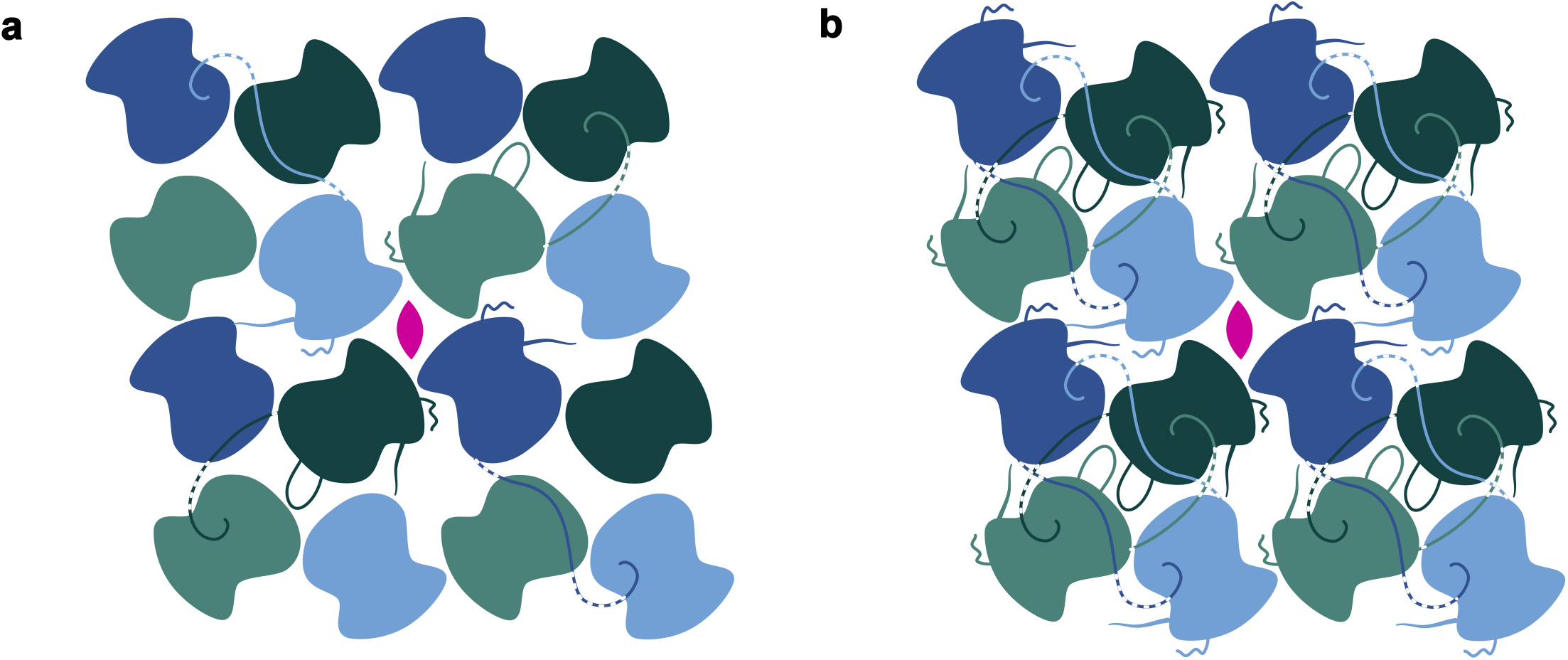
PhuN assembles through a series of complex and likely adaptive C-terminal tail exchanges. **a**, Cartoon showing core tetramer with N- and C-terminal tail interactions as well as the loops visible clearly in 2D classes. **b**, Cartoon model tracing the N- and C-terminal tail interactions at all four unique interfaces.

While we do not observe the N-terminal MBP tag in our maps, the MBP appears on the exterior of our small *in vitro* assemblies **(Fig. S1d)**. From this, we suggest that the lattice surface with the exposed N-terminus faces the cytosol. Thus, the exterior of the Phage Nuclear Shell is sporadically positively charged while the interior surface is quite negatively charged **(Fig. 3b)**. This could be an elegant strategy for keeping the DNA away from the surface, preventing it from inadvertently leaving the protection of the lattice network.

## Discussion

Once thought to be a defining feature of eukaryotic organisms, the Phage Nucleus proves to be a protein-based, membrane-free structure accomplishing a similar task. In this work we describe the novel self-assembly of the 66.6 kDa PhuN protein from bacteriophage ϕPA3 into a C2 symmetric lattice that closely matches fragments of the endogenous Phage Nucleus isolated from ϕPA3 infected *P. aeruginosa*. This tetrameric lattice features four distinct interfaces, two of which form clear channels (Diamond, Open Hairpin) that may allow the passage of small molecules, unfolded proteins, or nucleotides. We further show that the lattice assembles through interactions between flexible loops and establishes a negatively charged interior surface for the Phage Nucleus.

These lattices and channels provide a novel framework with which to approach further mechanistic questions in this system ranging from growth and adaptability of the lattice, to DNA protection and packaging, to selective translocation in and out of the Phage Nucleus. The channels we identify may allow for diffusion of small molecules as is or perhaps can accommodate larger molecules following conformational changes induced at specific sites or in response to binding of other phage factors. The different interactions of the large flexible loops and possibility for varied C-terminal tail binding observed here could help accommodate such changes. This idea is supported by a recent report of C4 symmetric lattices created by phage 201ϕ2-1.^17^ We also observed a minor C4 population in our 2D classes, suggesting that this symmetry may also be accessible in ϕPA3 PhuN assemblies **(Fig. S8a)**. The fact that the same monomer can readily form distinct PhuN symmetry states implies an unexpectedly flexible, dynamic assembly far different from the highly stereotyped, rigid lattices found in phage heads or compartments like the carboxysome.^10,11,25,26^ *In vivo*, scattered throughout this remarkable lattice may be other protein-based channels to allow for capsid docking, efficient DNA extrusion and packaging into empty capsids, as well as the passage of other large molecules.

In many other protein-based assemblies, such as eukaryotic and prokaryotic viral capsids and bacterial microcompartments, space is enclosed through combinations of triangular or hexametric units forming flat faces and pentameric units providing curvature.^10,11,26^ At the molecular level, the ability of the same proteins to pack in these different modes arises from quasi-equivalent interactions amongst flexible domains.^26^ Long tails are often used around the symmetry points to stabilize the assemblies.^26^ The observations reported here suggest that quasi-equivalence may also be mediated directly by flexible tails and loops, not only forming quasi-symmetric tetramers but also flexibly linking adjacent tetramers together.

Given the extensive interactions between tetramers mediated by these long tails and loops, adding new subunits during growth while maintaining a permeability barrier may require the transient formation of dislocations or perhaps the aid of helper host proteins, such as chaperones, as with clathrin lattices.^27^ The observation of small assemblies, similar to those found *in vitro*, attached to an isolated shell **(Fig. S1a)** also raises the possibility of a Phage Nuclear shell growth mechanism driven by the incorporation of small, pre-formed assemblies rather than individual monomers. The regular spacing and close proximity of these assemblies may also point to the existence of specialized regions along the Phage Nucleus where subunit addition occurs. This could allow for rapid and efficient growth while minimizing the number of disruptions made to the Phage Nuclear Shell lattice. Alternatively, these small assemblies may correspond to distinct protein complexes with quite different functional roles.

The flexibility of the C-terminal tail, availability of some variation in its binding to neighboring subunits, and potential for toggling between symmetric states is perhaps what enables the astounding diversity in both shape and size of the bacteriophage nuclei,^4^ especially considering they appear to be largely constructed from a single ∼70 kDa protein across various jumbo phages.^5^ Notably, PhuN proteins from different species do not appear to interact during co-infections,^28^ suggesting that diversity in the tails may contribute to phage speciation. As differing N- and C-terminal tail lengths account for the primary structural differences between AlphaFold predictions of known PhuN proteins **(Fig. S9)**, a closer and careful analysis of PhuN evolution from diverse jumbo bacteriophages will be instrumental in delineating mechanistic similarities and differences that underly the ability of PhuN proteins to assemble into the micron-scale Phage Nuclei observed *in vivo*. Honing in on these questions will provide a more thorough understanding of where these structures came from, how they nucleate and grow, as well as how we can co-opt them for both experimental and practical applications going forward.

## Acknowledgements

We thank past and present members of the Agard laboratory for many helpful discussions and support throughout this project; particularly, Feng Wang for functionalized EM grids and Shawn Zheng for developing *Motioncor2*; David Bulkley, Eric Tse, Glen Gilbert (W.M. Keck Foundation Advanced Microscopy Laboratory at UCSF, Mission Bay), and Alexander Myasnikov (current: Saint Jude) for maintaining the cryoEM facility and assistance with data collection; Matt Harrington and Joshua Baker-LePain for computational support with the UCSF Wynton Cluster; Amanda Drennan (Rayment Laboratory, UWMadison) for teaching E.S.N. how to prepare 2D crystals. Molecular graphics and analyses performed with UCSF Chimera (NIH P41-GM103311) and UCSF ChimeraX (NIH R01-GM129325); This research was supported by: NIH grant R35GM118099 (D.A.A.) and NIH facilities grants 1S10OD026881 (D.A.A.), 1S10OD020054 (D.A.A.), 1S10OD021741 (D.A.A.), Microsoft (M.B., D.B.), Open Philanthropy and HHMI (D.B.), the Washington Research Foundation (M.B.), NIH grant R01GM127489 (J.B.-D.), UCSF Program for Breakthrough Biomedical Research funded in part by the Sandler Foundation (J.B.-D.), NIH R35GM140847 and HHMI (Y.C.).

## Author Contributions

ESN performed cloning, experimental design, *in vitro* purification, sample preparation, EM data collection and analysis, model fitting, as well as coordinated experimental design for fragment isolations with MM, and trained MM in negative stain EM all under the supervision of DAA. AFB conducted motion correction of collected cryoEM data and advised EM processing, experimental design, and interpretation. MM isolated shell fragments. MB and DB provided early access to RoseTTAFold. JL and YC EM conducted EM map deconvolution and resolution estimation. CK and JBD contributed to editing the manuscript and intellectual discussion.

## Competing Interests

The authors declare no competing interests.

**Correspondence and requests for materials** should be addressed to David A. Agard.

