## SupplementaryMaterials for "The ϕPA3 Phage Nucleus is Enclosed by a Self-Assembling, 2D Crystalline Lattice"

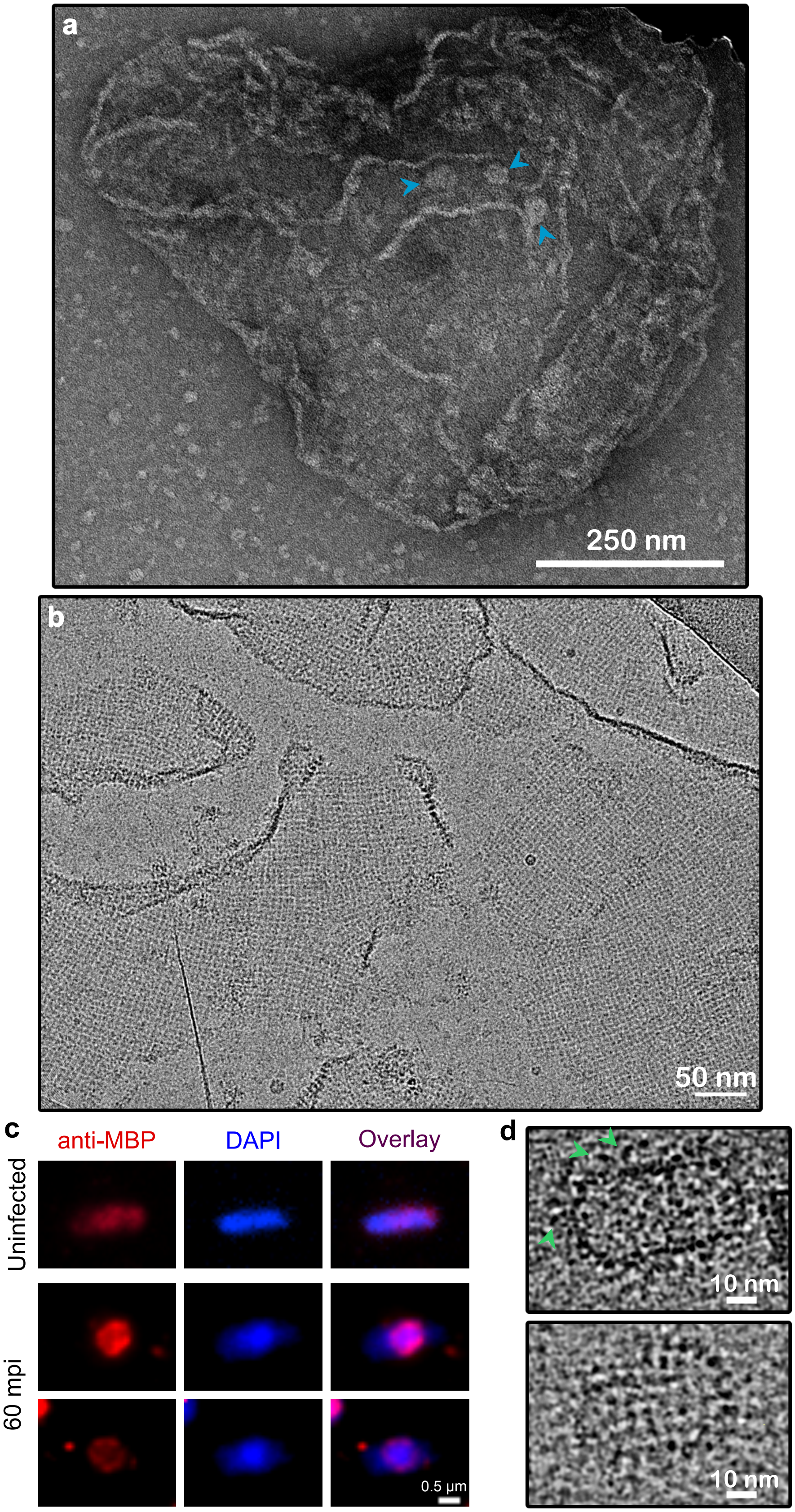


**Extended Data Fig. 1 | ϕPA3 PhuN assemblies. a,** Negative Stain micrograph of isolated Phage Nuclear Shell fragment about the size expected for an intact compartment. Lattice texture is visible as well as many wrinkles and folds. Smaller, round species are visible on the surface (blue arrows), similar in appearance to Class II assemblies *in vitro*. **b,** Full micrograph from Figure 1a showing many shell fragments with a lattice texture. **c,** Immunofluorescence microscopy of *P. aeruginosa* cells expressing the 6XHisMBPGp53 fusion. The fusion protein integrates into the shell and localizes at the cell center around the phage DNA 60 minutes post infection. **d,** CryoEM images of *in vitro* PhuN assemblies. MBP is visible on the exterior (green arrow, top image). The assemblies show a lattice like surface (bottom).

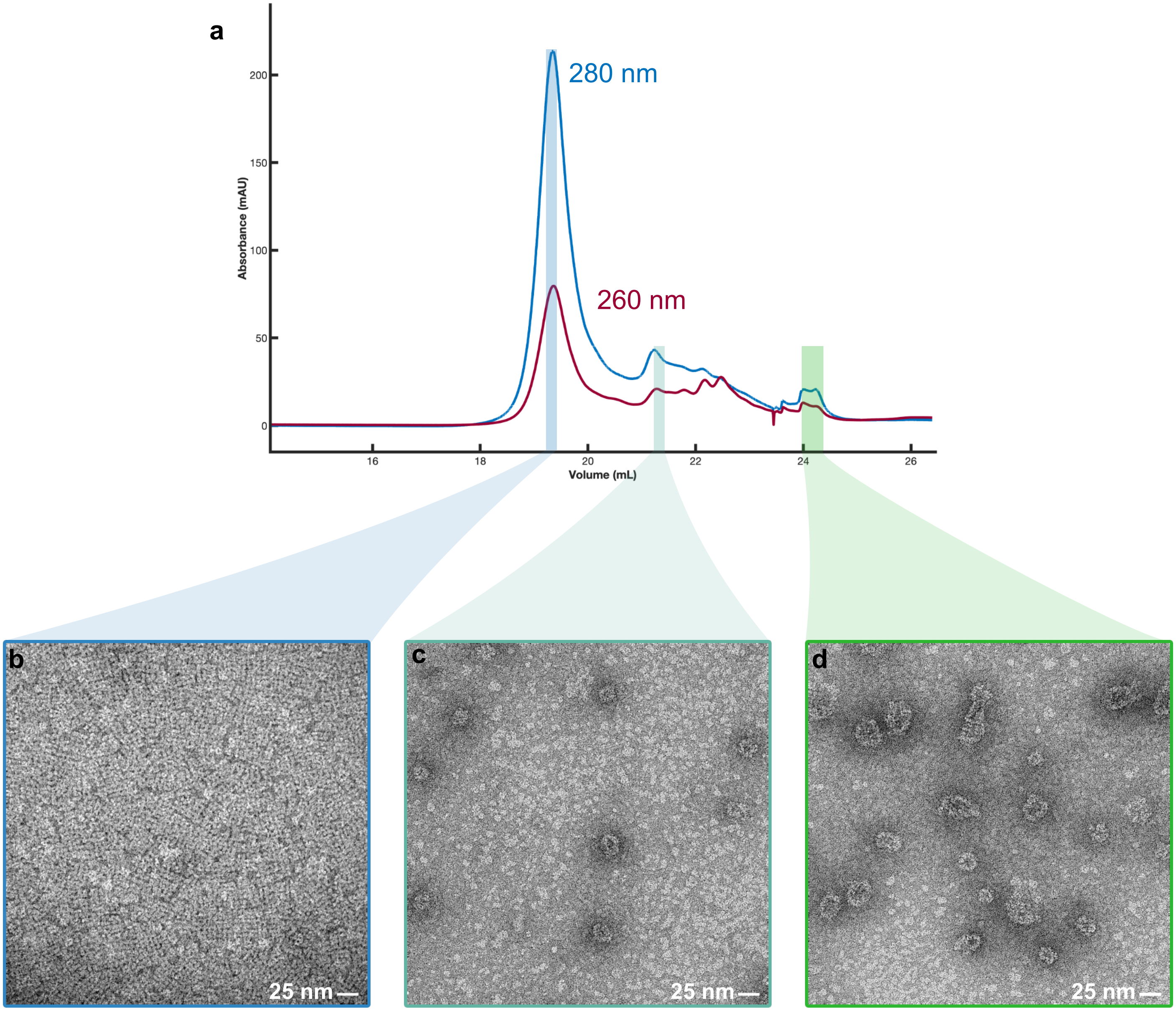


**Extended Data Fig. 2 | PhuN Crystalizes at pH 6.5. a.** Anion exchange purification using a 1 mL GE MonoQ column at pH 6.5. Highlighted fractions correspond to micrographs b-d. **b,** Monomeric species in Class I form 2D polycrystalline assemblies when applied to negatively charged grids for negative stain EM. **c,** 25 nm mid-sized species present in Class II. **d,** Assemblies 25 nm and larger present in later elution in Class III.

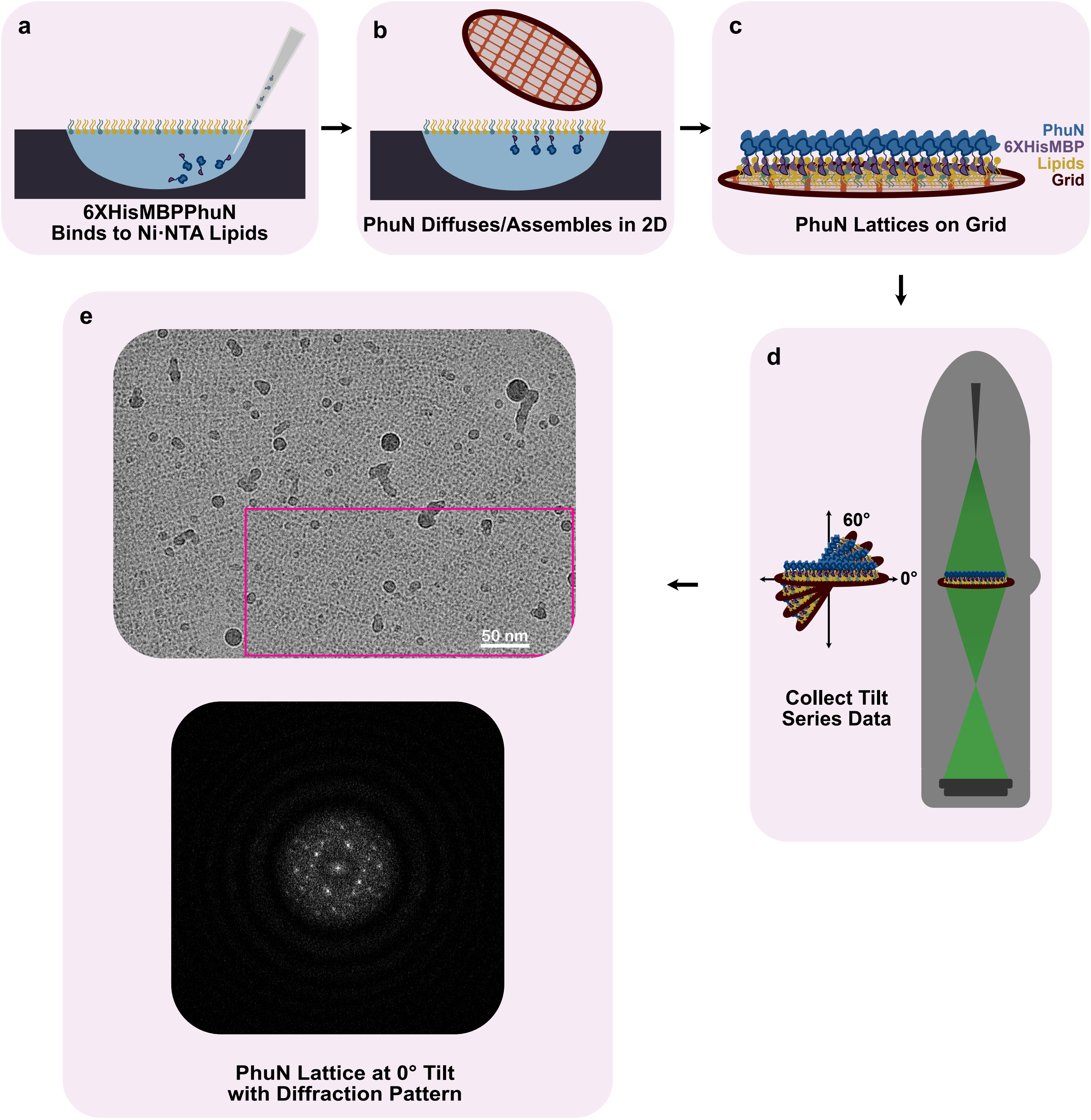


**Extended Data Fig. 3 | *in vitro* 2D Crystal Preparation. a,** 6XHisMBP-PhuN is injected into a buffer droplet with a pre-formed 21% Ni·NTA Lipid Monolayer. **b-c,** After incubation to allow for assembly, the assembled arrays are adsorbed on a grid and plunge frozen for CryoEM. **d,** Data is collected at fixed tilts ranging from 0-60°. **e,** 0° image from Fig. 1c with the diffraction pattern corresponding the region denoted with the magenta rectangle.

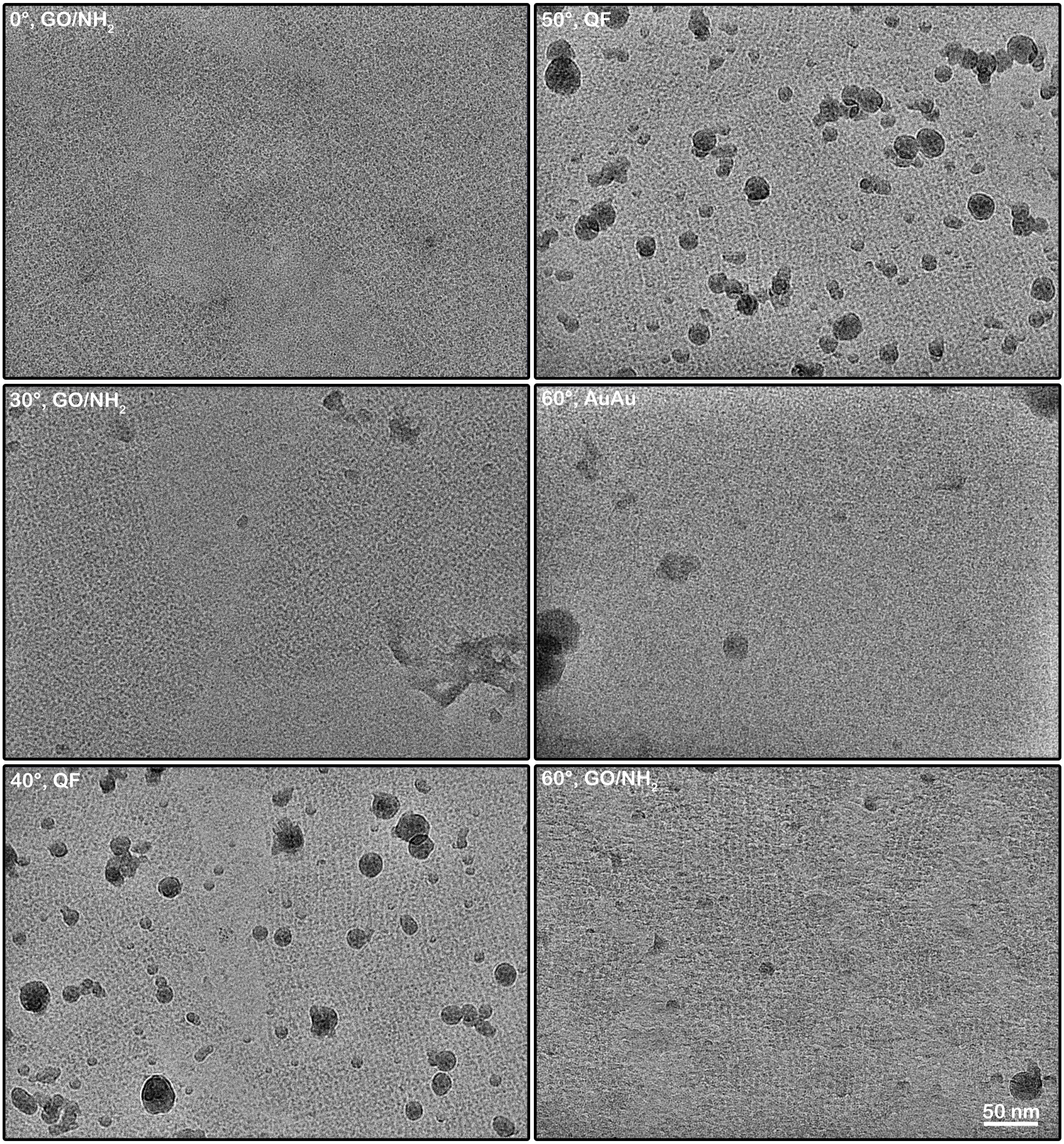


**Extended Data Fig. 4 | Example micrographs collected at the specified tilt angle.** Tilt angle is listed alongside the grid support used in each micrograph: Traditional Quantifoil (QF), Amino-functionalized graphene-oxide (GO/$NH_2$), UltrAuFoil (AuAu). The lattices often appear with bends and breaks as seen throughout these micrographs.

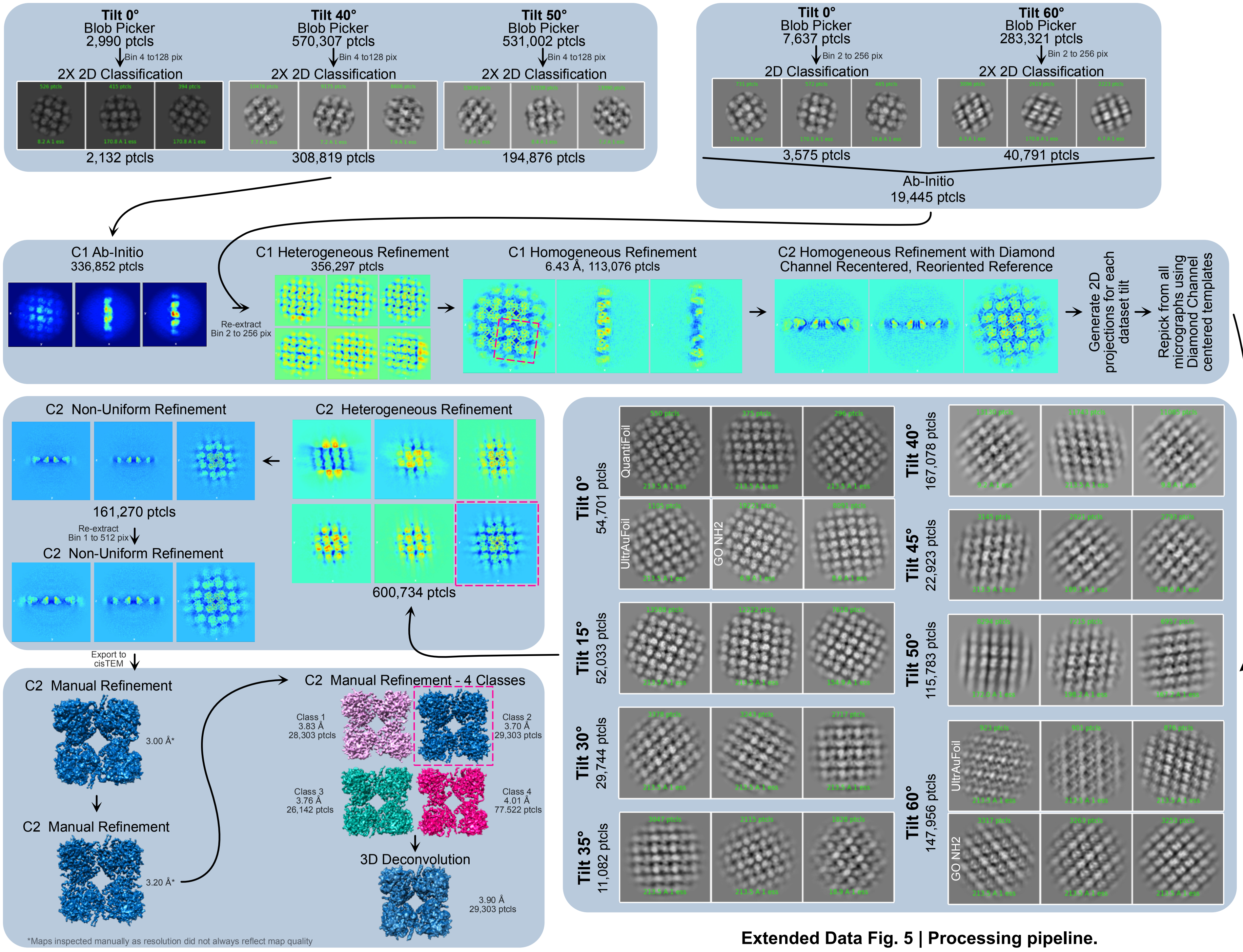


**Extended Data Fig. 5 | Processing pipeline.**

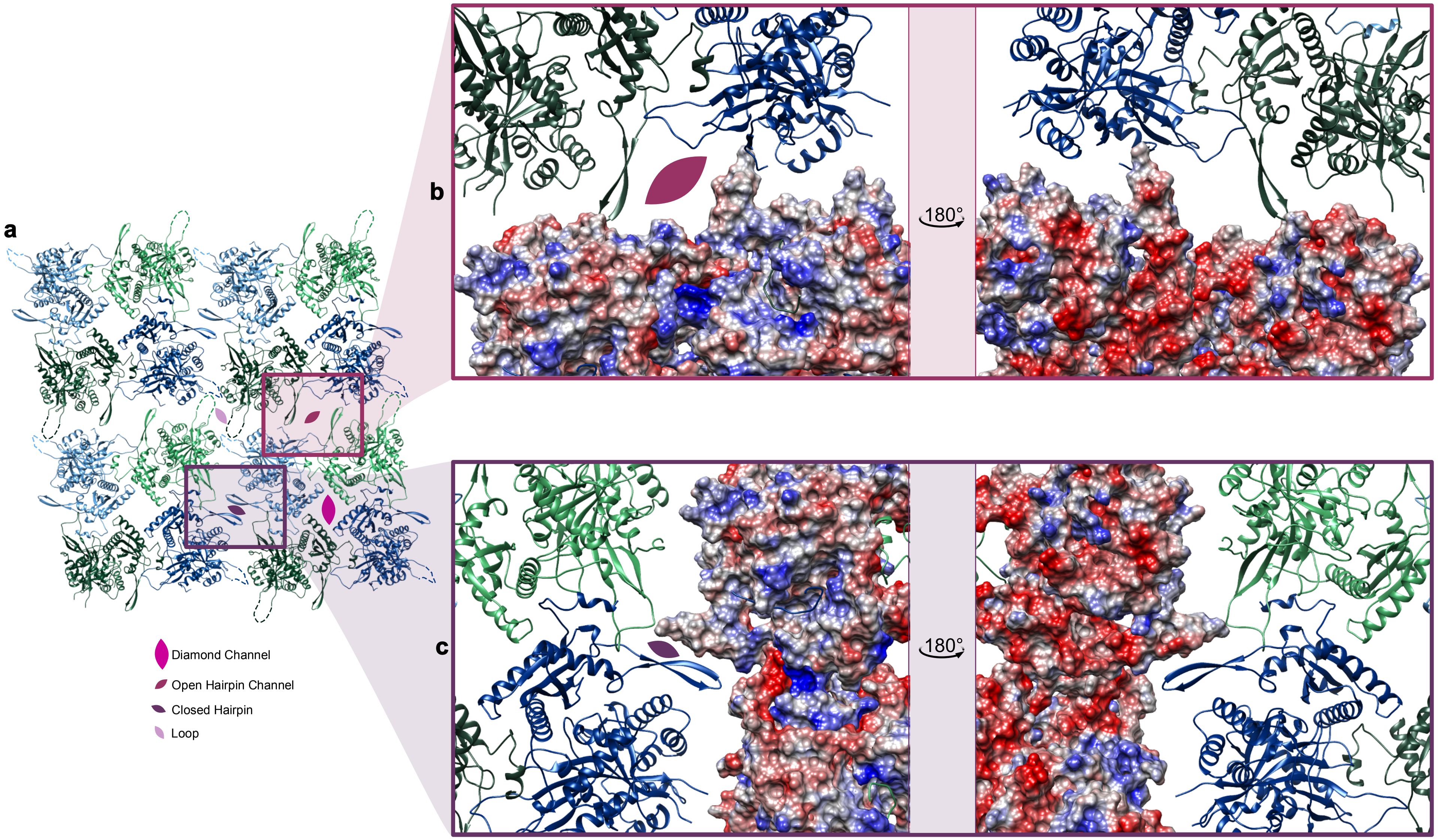


**Extended Data Fig. 6 | Closer look at the positioning of the β-Hairpin at the Open Channel and Closed Interface. a,** 16-mer model as in Fig. 2e. The two channels and two interfaces are labeled. **b,** The Open Channel is shown as viewed from the cytosol (left) and inside the Phage Nucleus (right). **c,** The Closed interface is similarly represented with the Phage Nucleus exterior on the left and interior on the right. The surfaces of two of the four subunits at each interface are shown and colored by electrostatic potential.

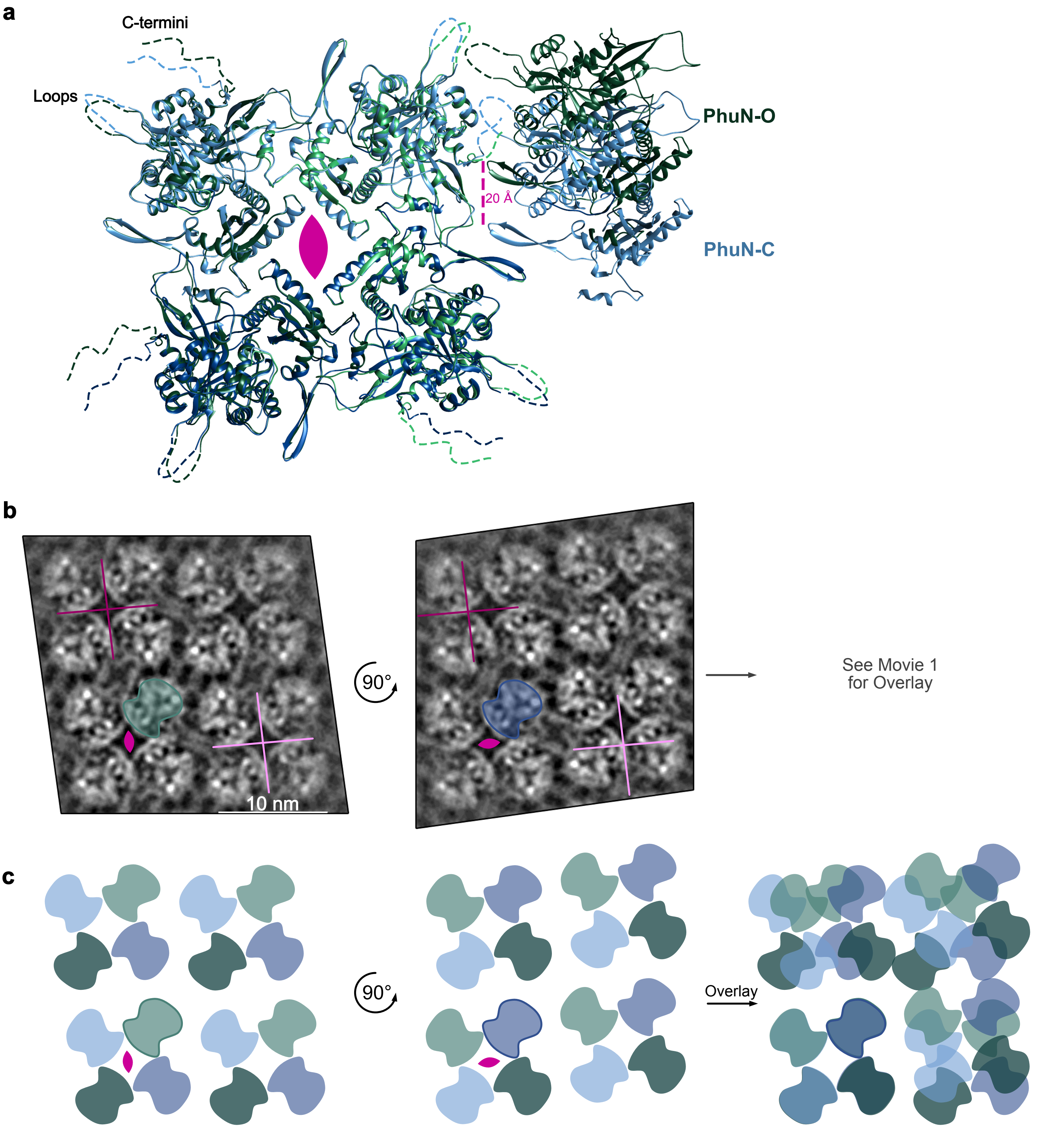


**Extended Data Fig. 7 | Comparison of the Open and Closed β-Hairpin Interfaces. a,** Two tetramers aligned on the asymmetric subunits with a neighboring PhuN-O and PhuN-C subunit. The β-hairpin of Phun-O is near the C-terminus of its neighbor while the PhuN-C β-hairpin is closer to a structured loop. **b,** 2D class, as shown in figure 2a, compared to a 90° rotation of the same 2D class after alignment on the highlighted asymmetric PhuN-O (green) and PhuN-C (blue) subunits. The centers of two Diamond Channels are shown with a light and dark pink cross (left) and are kept in the same position in the rotated image (right) to show the ~20 Å lattice shift. See Movie1 for a morph between the aligned subunits. **c,** Cartoon depiction of panel (a) with a direct overlay of the rotated lattice showing the new positioning of the shifted subunits. The alignment was done using the bolded PhuN-O (green) and PhuN-C (blue) as in panel (b).

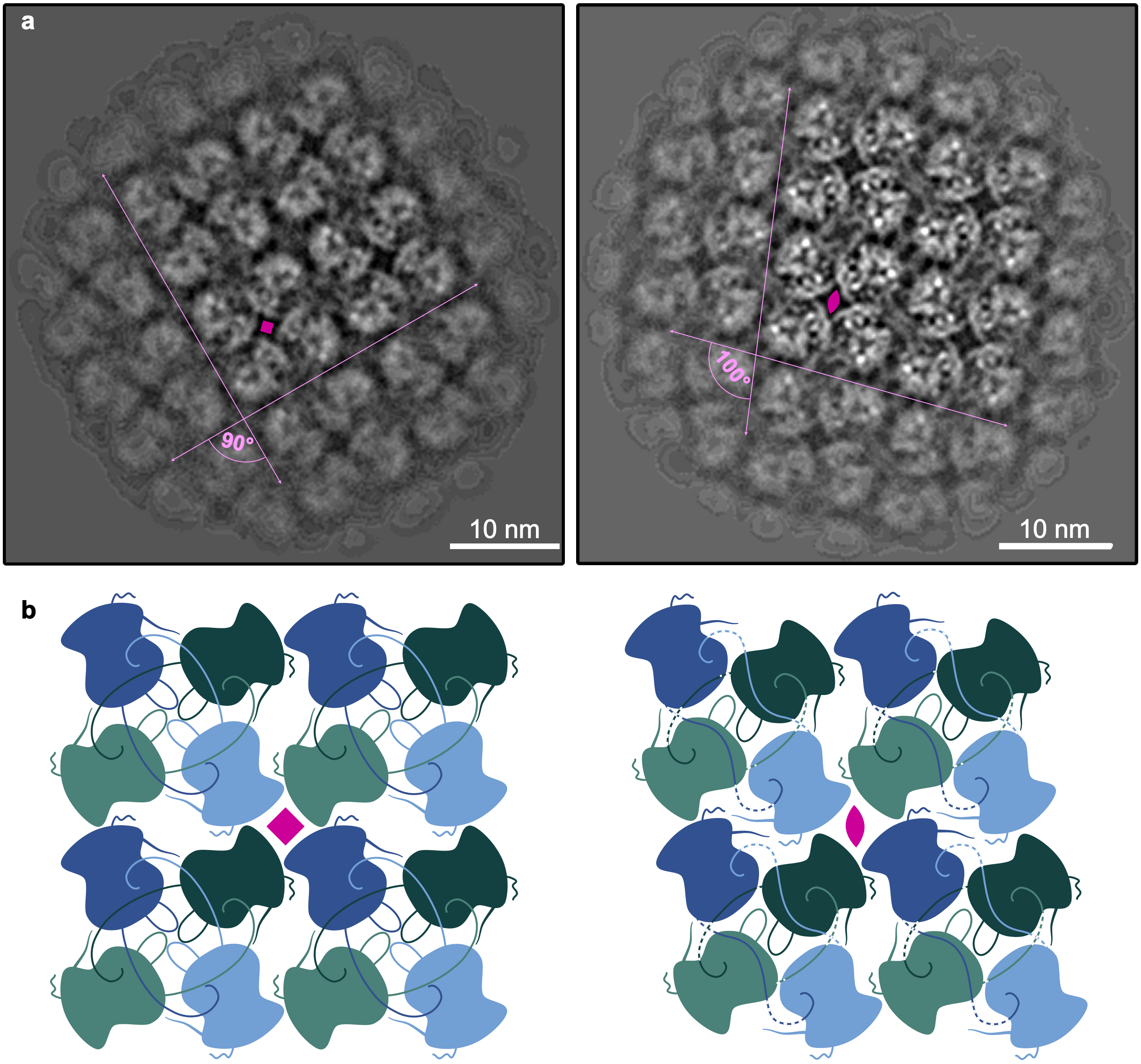


**Extended Data Fig. 8 | Comparison of C2 and C4 lattice symmetries. a,** A small population with C4 symmetry (left) is observed in 2D classification. The flexible loops at the Loop interface appear to all face the center while the remaining two interfaces are replaced by one that is most similar to the Open Hairpin interface in our C2 classes (right, full view of the class shown in Fig. 2a). **b,** Cartoon depiction of how the C4 and C2 lattices may compare

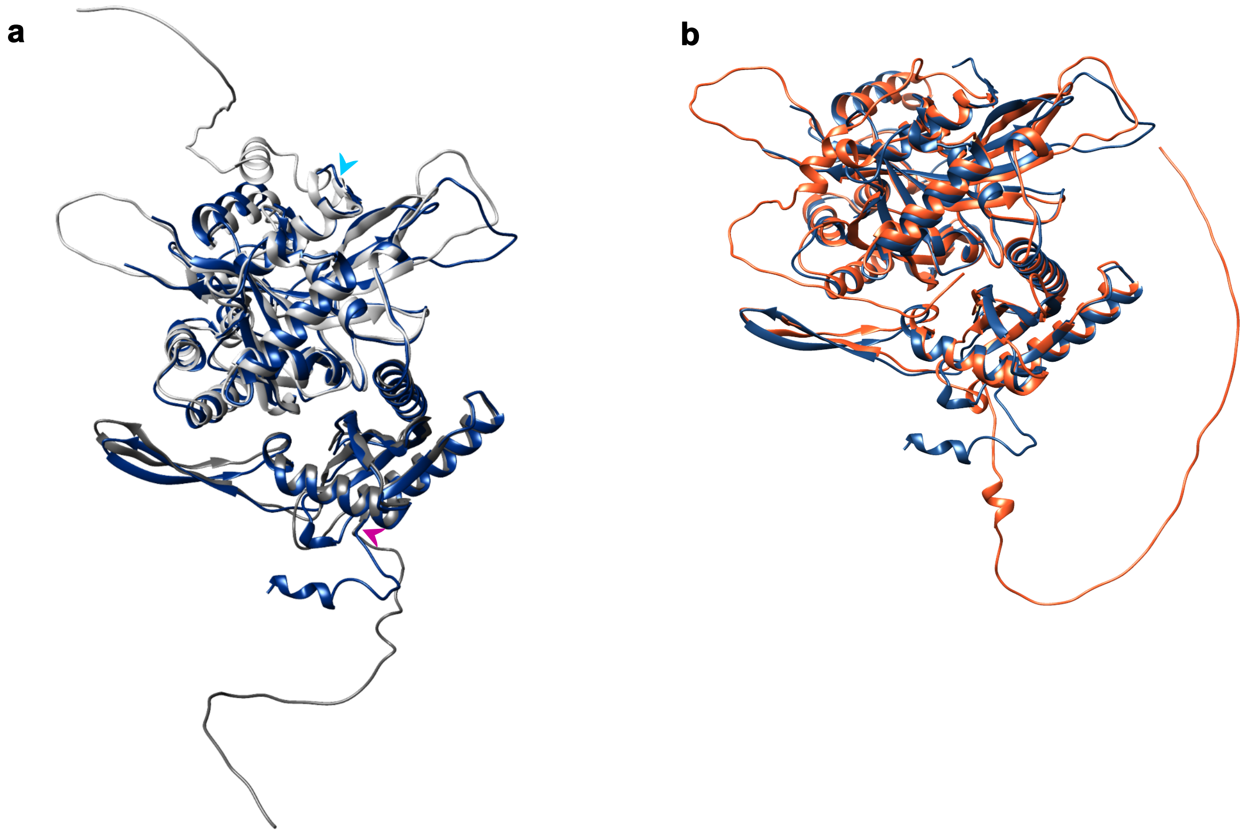

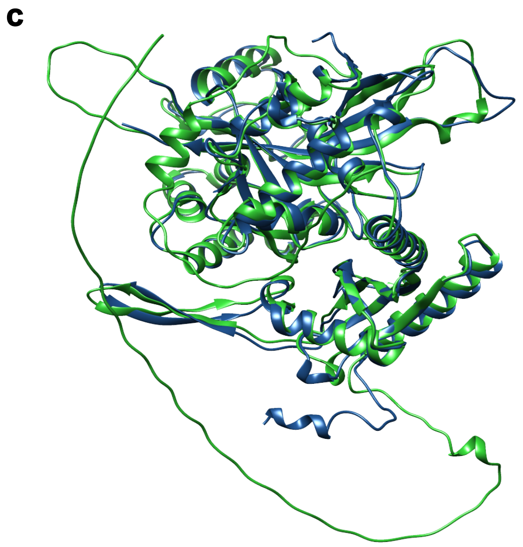

| d | ϕPA3 | ϕKZ | 201ϕ2-1 |
|---|---|---|---|
| **N-terminal Residues** | 36 | 53 | 64 |
| **C-terminal Residues** | 50 | 69 | 50 |

**Extended Data Fig. 9 | AlphaFold PhuN predictions compared to the final ϕPA3 PhuN model.** Final model of ϕPA3 gp53 (dark blue) overlaid with **a,** the starting ϕPA3 AlphaFold Prediction (grey), **b,** ϕKZ AlphaFold prediction (orange), and **c,** the 201ϕ2-1 AlphaFold prediction (green). In (b) and (c), the C-terminal tails are positioned across the back of the large, acetyltransferase-like domain. **d,** A summary of differing N- and C-terminal tail lengths as measured to the start of the helix lining the Diamond channel (pink arrow in panel a) for the N-term and from the end of the β-strand for the C-term (turquoise arrow in panel a).

| NTD | | | | | | |
|---|---|---|---|---|---|---|
| PDB ID-Chain | Z | rmsd | lali | nres | %ID | PDB Description |
| 3i01-C | 5.8 | 3.8 | 103 | 673 | 3 | CARBON MONOXIDE DEHYDROGENASE/ACETYL-COA SYNTHASE |
| 3i39-X | 5.7 | 3.9 | 99 | 635 | 9 | CARBON MONOXIDE DEHYDROGENASE 2 |
| 1su6-A | 5.7 | 4.0 | 102 | 633 | 8 | CARBON MONOXIDE DEHYDROGENASE 2 |
| 6x5k-C | 5.7 | 3.8 | 102 | 672 | 3 | CARBON MONOXIDE DEHYDROGENASE/ACETYL-COA SYNTHASE |
| 3nat-A | 5.6 | 3.1 | 93 | 156 | 17 | UNCHARACTERIZED PROTEIN |
| 2z8y-A | 5.6 | 3.5 | 99 | 673 | 3 | CARBON MONOXIDE DEHYDROGENASE/ACETYL COA SYNTHASE |
| 7b70-J | 5.6 | 4.5 | 111 | 614 | 6 | TRAFFICKING PROTEIN PARTICLE COMPLEX SUBUNIT |
| 3s1s-A | 5.5 | 3.6 | 100 | 871 | 7 | RESTRICTION ENDONUCLEASE BPUSI |
| 1su7-A | 5.5 | 3.7 | 99 | 633 | 10 | CARBON MONOXIDE DEHYDROGENASE 2 |
| 2yiv-X | 5.5 | 3.9 | 98 | 635 | 8 | CARBON MONOXIDEDEHYDROGENASE 2 |
| 5hci-F | 5.5 | 3.5 | 96 | 249 | 4 | GPN-LOOP GTPASE1 |
| 7b6z-A | 5.5 | 4.5 | 111 | 614 | 6 | TRAFFICKING PROTEIN PARTICLE COMPLEX SUBUNIT 11 |
| 3hrl-A | 5.4 | 3.3 | 90 | 94 | 4 | ENDONUCLEASE-LIKE PROTEIN |
| 7p2q-C | 5.4 | 3.7 | 85 | 174 | 11 | SIGNAL PEPTIDASE COMPLEX CATALYTIC SUBUNIT SEC11C |
| 4udy-X | 5.4 | 3.9 | 100 | 633 | 7 | CARBON MONOXIDE DEHYDROGENASE 2 |
| 1mjg-B | 5.4 | 3.8 | 102 | 672 | 3 | CARBON MONOXIDE DEHYDROGENASE BETA SUBUNIT |
| 3i01-A | 5.3 | 3.5 | 101 | 673 | 3 | CARBON MONOXIDE DEHYDROGENASE/ACETYL-COA SYNTHASE |
| 1oao-B | 5.3 | 3.8 | 102 | 673 | 3 | CARBON MONOXIDE DEHYDROGENASE/ACETYL-COASYNTHASE |
| 5hci-E | 5.3 | 3.7 | 101 | 259 | 6 | GPN-LOOP GTPASE 1 |
| 1oao-A | 5.2 | 3.9 | 103 | 673 | 3 | CARBON MONOXIDE DEHYDROGENASE/ACETYL-COA SYNTHASE |

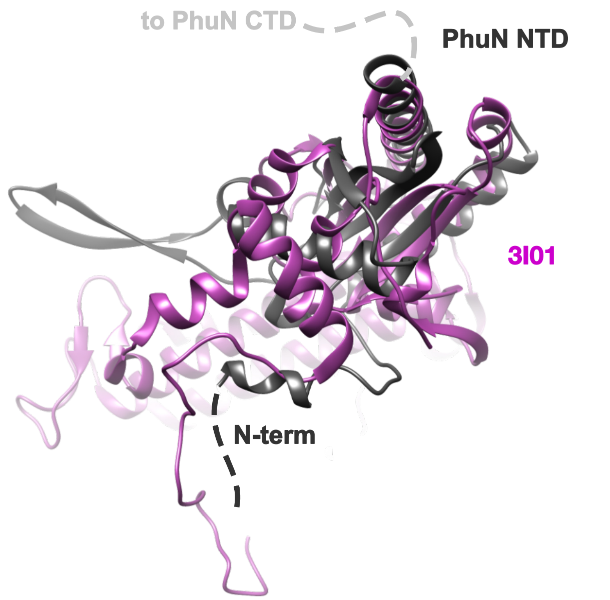


| CTD | | | | | | |
|---|---|---|---|---|---|---|
| PDB ID-Chain | Dali Z-score | rmsd | lali | nres | %id | PDB Description |
| 7byy-A | 6.0 | 3.9 | 130 | 180 | 10 | ACETYLTRANSFERASE |
| 7ak9-B | 5.9 | 3.7 | 126 | 163 | 11 | ACETYLTRANSFERASE |
| 7ak9-A | 5.8 | 3.7 | 125 | 162 | 10 | ACETYLTRANSFERASE |
| 7byy-C | 5.8 | 4.0 | 129 | 176 | 9 | ACETYLTRANSFERASE |
| 6g96-B | 5.7 | 4.1 | 129 | 174 | 10 | ACETYLTRANSFERASE |
| 6g96-A | 5.6 | 3.9 | 125 | 175 | 10 | ACETYLTRANSFERASE |
| 7chd-C | 5.6 | 4.0 | 130 | 179 | 12 | N-ACETYLTRANSFERASE DOMAIN-CONTAINING PROTEIN |
| 7byy-D | 5.5 | 4.1 | 124 | 173 | 9 | ACETYLTRANSFERASE |
| 7byy-B | 5.5 | 4.2 | 128 | 179 | 9 | ACETYLTRANSFERASE |
| 6cw2-D | 5.4 | 3.7 | 129 | 241 | 7 | HISTONE ACETYLTRANSFERASE GCN5 |
| 3f5b-A | 5.4 | 3.2 | 122 | 172 | 5 | AMINOGLYCOSIDE N(6')ACETYLTRANSFERASE |
| 2qml-A | 5.4 | 3.3 | 126 | 193 | 10 | BH2621 PROTEIN |
| 1y9k-B | 5.4 | 3.7 | 120 | 154 | 8 | IAA ACETYLTRANSFERASE |
| 6ajm-A | 5.4 | 4.2 | 127 | 172 | 10 | N-ACETYLTRANSFERASE |
| 6ajn-A | 5.4 | 4.3 | 126 | 172 | 10 | N-ACETYLTRANSFERASE |
| 7chd-B | 5.3 | 4.1 | 128 | 180 | 11 | N-ACETYLTRANSFERASE DOMAIN-CONTAINING PROTEIN |
| 6gtq-A | 5.3 | 4.0 | 122 | 160 | 11 | N-ACETYLTRANSFERASE |
| 6ajm-B | 5.3 | 3.6 | 118 | 157 | 10 | N-ACETYLTRANSFERASE |
| 5w8e-A | 5.2 | 3.5 | 127 | 214 | 4 | AUTOINDUCER SYNTHASE |
| 6gtr-A | 5.2 | 4.2 | 121 | 160 | 12 | N-ACETYLTRANSFERASE |

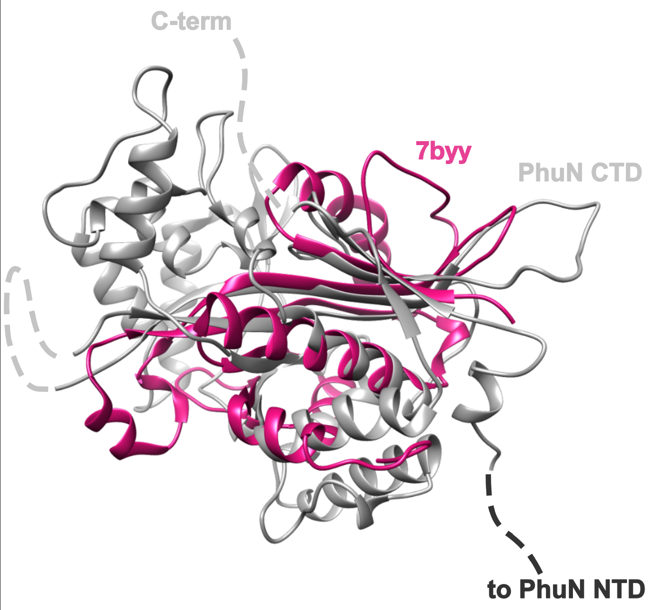


**Extended Data Fig. 10 | Dali Server searches of the PhuN NTD and CTD.** 20 highest scoring Dali Server outputs with the NTD (top) and CTD (bottom) as the input. The highest scoring structural homologue is shown to the right of the corresponding table.

### Methods

*6XHisMBPGP53 Construct and Expression*
An N-terminal maltose binding protein (MBP) tag and *E. coli* codon optimized ϕPA3 gp53 gene were inserted into a pET15 backbone to create pESN6. pESN6 was transformed into BL21(DE3) Star cells for expression. 20 mL of an overnight in LB was grown with carbenicillin and chloramphenicol. 1L of Terrific Broth (TB) growth media was inoculated with around 2% of the overnight or until the starting $OD_{600}$ reached 0.001. Cells were grown with 250 rpm nutation at 37°C until an $OD_{600}$ of 0.5. Cells and the incubator were then cooled to 16°C. Expression was induced with 1 mM IPTG and proceeded overnight at 16°C with 250 rpm nutation. Cells were spun down the following morning at 3,000 rcf for 8 minutes and pellets were transferred to 50 mL conical tubes. These were flash frozen and stored at -80°C or used immediately for purification.

*Phage Shell Isolation*
*P. aeruginosa* PA01 expressing MBP-gp53 were lysed at 60 min post ϕPA3 infection in NP40 Lysis Buffer (50 mM Bis-Tris, 150 mM NaCl, 0.5% NP40, 5% glycerol, 5 mM DTT, 20 ng/µl Lysozyme, 1 mM EDTA, 1mM EGTA – pH 6.5) by dounce homogenizer. The lysate was clarified at 16000xg for 5 min and the insoluble fraction resuspended in Wash/Shell buffer (20 mM Bis-Tris, 150 mM NaCl, 1 mM DTT, 1 mM EDTA, 2 mM $MgCl_2$ – pH 6.5). This was subject to further 500xg (5 min) and 15000xg (10 min) centrifugation with the insoluble fraction isolated and resuspended in Wash/shell buffer each time. The final 15000xg pellet resuspension was used directly for downstream analysis by negative stain or cryo-EM.

*Immunofluorescence Microscopy*
Immunofluorescence microscopy samples were prepared as previously published.[3]

*6XHisMBPGP53 Purification*
A cell pellet from 1 L of culture was resuspended in 12 mL of Lysis Buffer pH 7.42 (20 mM HEPES, 500 mM NaCl, 0.2% TritonX100, 1 mM DTT, 5% Glycerol, 5 mM $MgCl_2$) per each gram of pellet with 1 cOmplete Protease Inhibitor Tablet and benzonase. This mixture was homogenized using a manual glass homogenizer and lysed using an Avestin EmulsiflexC3. The sample was clarified with a 20 minute spin at 18,500 rcf. The soluble fraction was nutated with equilibrated NiNTA resin for 1-2 hours in the cold room. The sample was transferred to a gravity column and washed 2X with the ATP Wash Buffer pH 7.42 (5 mM ATP, 5 mM $MgCl_2$, 20 mM HEPES, 500 mM NaCl, 20 mM Imidazole, 1 mM DTT, 5% Glycerol) and 2X with Wash Buffer pH 7.42 (20 mM HEPES, 500 mM NaCl, 20 mM Imidazole, 1 mM DTT, 5% Glycerol). Finally, the sample was eluted and collected in the Elution Buffer pH 7.42 (20 mM HEPES, 150 mM NaCl, 600 mM Imidazole, 1 mM DTT, 5% Glycerol). The eluate was aliquoted and flash frozen the same day and stored at -80°C.

*Anion Exchange*
The sample (thawed or fresh) was buffer exchanged into MonoQ Binding Buffer (pH 7.65, 20 mM Tris, 50 mM NaCl, 1 mM DTT, 0.5 cOmplete Protease Inhibitor Tablet; pH 6.5, 20 mM Bis-TrisPropane, 50 mM NaCl, 1 mM DTT, 1 mM EDTA, 0.5 cOmplete Protease Inhibitor Tablet) using 3.5K MWCO Snakeskin dialysis tubing or 7K MWCO Zeba Spin Desalting Column. The sample was loaded on the 10/100 or 5/50 GL MonoQ Anion Exchange Column, washed, and eluted over a linear gradient from 0 to 65% over 13 mL (5/50 gL) using the Elution Buffer (pH 7.65, 20 mM Tris, 1 M NaCl, 1 mM DTT, 0.5 cOmplete Protease Inhibitor Tablet; pH 6.5, 20 mM Bis-TrisPropane, 1 M NaCl, 1 mM DTT, 1 mM EDTA, 0.5 cOmplete Protease Inhibitor Tablet)

*2D Crystal Preparation and Sample Freezing*

The most concentrated monomeric fraction was selected from the Anion Exchange purification. The concentration was used to calculate the volume of buffer and sample needed to reach a working concentration of 0.01 mg/mL 6XHisMBPgp53 in a 30 uL crystallization drop. A humidity chamber was made by wetting filter paper inside of a petri dish. A teflon block with approximately 5 mm wide and 13 mm deep wells was placed in this chamber. The calculated volume of 2D Crystallization Buffer (pH 6.5, 20 mM BisTrisPropane, 150 mM NaCl, 1 mM DTT, 1 mM EDTA, 2 mM $MgCl_2$, 1 mM ATP, 0.25 cOmplete Protease Inhibitor Tablet) was pipetted into each well. 1 $\mu$L of Lipid Mix (0.73 mg/mL EGG PC, 0.27 mg/mL DGS-NTA(Ni)) was dispensed onto each buffer droplet. The volume of concentrated protein was then gently added to each droplet. After a 1 hour incubation at room temperature, the lattices were adsorbed to amino-functionalized graphene oxide[22] or QUANTIFOIL R1.2/1.3 400-mesh copper holes carbon grids without glow-discharging. These were then plunge-frozen using a Vitrobot Mark IV (FEI) at 10°C with 100% humidity using a 4 second blot time, 0 second wait time, and blot force of 0 for the functionalized grids and a 6 second blot time, 0 second wait time, and blot force of 3 for the QUANTIFOIL grids.

*Acquisition of Cryo-Electron Microscopy Data*

As the data is composed of 15 collections varying tilt, grid type, and frame rates, the below description summarizes the ranges of settings for all of the collections combined.

The micrographs were collected using two 300 kV FEI Titan Krios electron microscopes equipped with K3 direct electron cameras (Gatan) and operated using SerialEM software. A nominal defocus range of 0.8-2.5 $\mu$m under focus was used for collection. A nominal magnification of 105 kx, corresponding to a physical pixel size of 0.835 Å or 0.834 Å, was used. A total dose of 66 or 67 e-/Å$^2$ was maintained for each collection. The samples were tilted from 0° to 60° for static dataset collection to obtain side-views of the 2D crystalline sample. To mitigate beam-induced motion, we utilized GO/$NH_2$ coated R1.2/1.3 300 mesh gold grids and collected data using a higher frame rate to obtain a finer sampling of the beam-induced motion.

| **Collected Datasets by Grid Type, Tilt, and Frame Rate** | | | | | | |
|---|---|---|---|---|---|---|
| **Grid Type** | Quantifoil | GO/$NH_2$ | GO/$NH_2$ | GO/$NH_2$ | UltrAuFoil | UltrAuFoil GO/$NH_2$ |
| **Tilt Angle** | 0°, 40°, 50° | 0°, 60° | 0°, 15° | 0°, 30°, 45°, 60° | 0°, 60° | 35° |
| **Frames** | 120 | 118 | 246 | 444 | 444 | 444 |
| **Magnification** | 105,000 | 105,000 | 105,000 | 105,000 | 105,000 | 105,000 |
| **Voltage (kV)** | 300 | 300 | 300 | 300 | 300 | 300 |
| **Electron Dose (e-/Å$^2$)** | 66 | 67 | 67 | 67 | 67 | 67 |
| **Defocus Range ($\mu$m)** | 1.5-2.5 | 0.8-1.6 | 0.8-1.6 | 0.8-1.6 | 0.8-1.6 | 0.8-1.6 |
| **Pixel Size (Å)** | 0.834 | 0.835 | 0.835 | 0.835 | 0.835 | 0.835 |
| **Combined Dataset Particle Processing** | | | | | | |

| | |
|---|---|
| **Symmetry Imposed** | C2 |
| **Number of Final Particles** | 29,303 |
| **Map resolution (Å)** | 3.8 |
| **FSC Threshold** | 0.143 |

*Processing Cryo-Electron Microscopy Data*

All micrographs were motion corrected using UCSF MotionCor2[21] with a 9 by 7 patch. Each tilt, frame rate, and grid type was imported and processed separately through 2D classification. The first datasets, collected on Quantifoil grids, were used to generate the first 3D model. After importing the motion corrected micrographs into cryoSPARC,[29] CTF estimation was done using cryoSPARC Patch CTF estimation (multi). cryoSPARC Blob Picker was then used to initially select particles. Two rounds of 2D classification were used to clean up picks. The selected particles from each tilt (0° 2,123 ptcls, 40° 308,891 ptcls, 50° 194,876 ptcls) were merged in a cryoSPARC Ab-Initio Reconstruction with C1 symmetry. A 176,652 particle class from the Ab-Initio Reconstruction gave a clear lattice volume with an apparent C2 symmetry. This volume was further used as a template for cryoSPARC Heterogenous Refinement.

Additional datasets underwent the same processing through 2D classification. Selected particles were added to the best particle subset for cryoSPARC Heterogenous Refinements with the best resolved volume as a template. The template was ultimately recentered on the diamond shaped channel at which point C2 symmetry was also imposed going forward. The centered template was also used to create a particle picking template for each tilted dataset. The template was generated using the relion project feature in RELION v.3.0.8,[38] saving a projection ±5° from the tilt at data collection every 3° and repeating this every 3° around the z axis from 0-180°. This yielded 244 templates for each tilted dataset. These templates were imported into cryoSPARC and all datasets were repicked using their respective template set. The picks were cleaned up with two rounds of 2D Classification resulting in ~600,000 particles. These better centered particles were classified using cryoSPARC Heterogenous Refinement and the resulting best class was processed using cryoSPARC Non-Uniform Refinement[30] to a reported 4.11 Å. These particles were exported for further processing in cisTEM.[39] In cisTEM, a mask was applied to the central tetramer for several rounds of Manual 3D Refinement.[39] The reported resolution estimates did not always correspond to changes in map quality, thus each map was visually inspected for improvement. The particles were further classified in 3D using cisTEM, resulting in a final class containing 29,303 particles.[39] This final volume was deconvolved to mitigate stretching along the z-axis due to the missing cone of information resulting from tilted data collection.

*Deconvolution of a map with preferred orientation*

Deconvolution was performed using ER-Decon II, which was initially developed to deconvolve 4D wide-field light microscopy volume with extremely low signal-to-noise ratio.[31] ER-Decon II is implemented in program PRIISM.[32] To apply ER-Decon II on a cryo-EM density map, an optical point spread function (PSF) is generated from the directional FSC of the cryo-EM map.[33] And the dFSC was used as a representation of preferred orientation. The smoothing and nonlinearity parameter for deconvolution was 0.5 and 10000, respectively. The detailed procedure and full evaluation of volume deconvolution will be described separately.

*Model Building*
RoseTTAFold[15] and AlphaFold,[16] were utilized to generate initial models. RoseTTAFold provided us with a first look at PhuN Gp53 while AlphaFold ultimately gave us a model which fit into the density slightly better, particularly after processing with RosettaRelax.[34] The N-terminus was positioned manually in ChimeraX Isolde[35] and further refined using RosettaRefine.[34] Residues 1-18, 276-287, and 556-602 did not have corresponding densities and were deleted for fitting. The model fitting was done using our deconvolved map, fitting each subunit within the tetramer separately. The best fitting asymmetric subunits from RosettaRefine[34] were selected and used to create a C2 symmetric tetrameric model. Invading C-terminal tails were similarly placed using Chimera, relaxed using ChimeraX Isolde,[35] and refined using RosettaRefine.[34] Structure validation was done using MolProbity.

*Data Analysis and Figure Preparation*
Figures were created using UCSF Chimera[36] and ChimeraX.[37] Anion exchange traces plotted using MatLab (MATLAB. (2021). *version 9.11.0. (R2021b).* Natick, Massachusetts: The MathWorks Inc.
